# Meta-exoproteomics identifies active plant-microbe interactions operating in the rhizosphere

**DOI:** 10.1101/2021.09.01.458574

**Authors:** Ian D.E.A. Lidbury, Sebastien Raguideau, Senlin Liu, Andrew R. J. Murphy, Richard Stark, Chiara Borsetto, Tandra Fraser, Andrew Goodall, Andrew Bottrill, Alex Jones, Gary D. Bending, Mark Tibbet, John P. Hammond, Chris Quince, David J. Scanlan, Jagroop Pandhal, Elizabeth M. H. Wellington

## Abstract

The advance of DNA sequencing technologies has drastically changed our perception of the complexity and structure of the plant microbiome and its role in augmenting plant health. By comparison, our ability to accurately identify the metabolically active fraction of soil microbiota and their specific functional role is relatively limited. Here, we combined our recently developed protein extraction method and an iterative bioinformatics pipeline to enable the capture and identification of extracellular proteins (meta-exoproteomics) expressed in the rhizosphere of *Brassica* spp. First, we validated our method in the laboratory by successfully identifying proteins related to the host plant (*Brassica rapa*) and a bacterial inoculant, *Pseudomonas putida* BIRD-1, revealing the latter expressed numerous rhizosphere specific proteins linked to the acquisition of plant-derived nutrients. Next, we analysed natural field-soil microbial communities associated with *Brassica napus* L (Oil Seed rape). By combining deep-sequencing metagenomics with meta-exoproteomics, a total of 1882 proteins were identified in bulk and rhizosphere samples. Importantly, meta-exoproteomics identified a clear shift (p<0.001) in the metabolically active fraction of the soil microbiota responding to the presence of *B. napus* roots that was not apparent in the composition of the total microbial community (metagenome). This metabolic shift was associated with the stimulation of rhizosphere-specialised bacteria, such as *Gammaproteobacteria*, *Betaproteobacteria* and *Flavobacteriia* and the upregulation of plant beneficial functions related to phosphorus and nitrogen mineralisation. By providing the first meta-proteomic level assessment of the ‘active’ plant microbiome at the field-scale, this study demonstrates the importance of moving past a genomic assessment of the plant microbiome in order to determine ecologically important plant: microbe interactions driving plant growth.

## Introduction

The plant microbiome is integral to plant health, as it delivers several life support functions^1, 2^. This includes enhancing the plant’s ability to acquire both macro- and micronutrients, such as nitrogen, phosphorus, and iron, as well as enhancing plant innate immunity against a range of plant pathogens^2–4^. Since the green revolution, intensive agricultural practices have resulted in a decoupling between microbes and their host plants^5^. The breakdown of rhizobia-legume symbiosis in heavily fertilised cropping systems is perhaps the most well-known example^6^. Others, such as the apparent reduction in the relative abundance of Bacteroidetes in domesticated crops relative to their wild cultivars, are more cryptic^7^. Agriculture is now facing a significant global crisis: a rapidly changing climate, an ever-growing human population, and depletion of our natural resources used to fuel crop production has identified severe vulnerabilities in ensuring future food security^2, 8^. Whilst the industrial production of nitrogen fertilisers is a highly energetic process, the production of inorganic phosphorus fertilisers is reliant on the continued supply of mined rock phosphate^9^. The latter of these fertiliser production regimes is set to cause various socioeconomic and political tensions as global stocks of rock phosphate are depleted^9, 10^. Thus, there is an urgent need to develop a holistic understanding of the plant microbiome function and its numerous components^11^.

Through the release of signalling molecules, exudation of organic nutrients and the decoration of plant cell walls with specific attachment molecules, plants can actively select for a subset of specialised soil microorganisms^12, 13^. This frequently involves a reduction in microbial diversity as one moves from the bulk soil > rhizosphere > root tissue^1, 14^. Whilst bulk soil is considered a relatively carbon poor environment favouring an oligotrophic lifestyle, the rhizosphere and root system is typified by a high turnover of organic matter driven through rhizodeposition, an environment favouring a copiotrophic lifestyle. Indeed, copiotrophic bacteria related to *Proteobacteria*, *Bacteroidetes* and *Actinobacteria* often dominate plant-associated microbial communities^15, 16^.

Whilst our understanding of the diversity, structure and functional potential of microbial communities has drastically improved, there is still considerable uncertainty about how this translates into specific plant: microbe interactions, especially carbon for nutrient exchange^2^. Therefore, we still lack understanding of the functional components involved in delivering beneficial plant activities within the root microbiome. Proteins are the functional entities of the cell whose regulation is controlled by surrounding biotic and abiotic conditions. Metaproteomics, the study of the entire protein content of a given environmental sample, holds enormous potential to improve our understanding regarding the function of soil microbial communities^17^. Unlike its application in seawater^18, 19^, anaerobic digestors^20, 21^ or the human or animal gut^22, 23^, soil metaproteomics is still in its infancy and conventional methods for the extraction of protein from soil are often plagued by the presence of contaminating substances, such as organic carbon and humic acids^24^. Furthermore, the majority of expressed proteins are related to cytoplasmic housekeeping and core metabolic functions, which can often result in poor detection of more ecologically importantly but less abundant non-cytoplasmic proteins^25^. One alternative is to focus on the extracellular (exo) fractions of proteins found outside the cell using meta-exoproteomics, a method which adapts extraction protocols for detecting soil extracellular enzyme activity^26^. Meta-exoproteomics has been successfully utilised to determine the active chitin degrading community of a tropical soil in response to chitin amendment^26^. Whilst this extraction method is applicable to bulk soil analysis (requiring 50-100 g soil material), sampling the rhizosphere (typically 1-2 g material) is much more challenging. Furthermore, the method currently requires specialised equipment and is relatively low throughput. These technical limitations have likely reduced the take up of this approach, despite its enormous potential.

Our recent work has successfully characterised the *in vitro* exoproteomes of single strain cultures related to *Pseudomonas* spp. and *Flavobacterium* spp. in response to phosphate-limiting growth conditions^27, 28^. These rhizobacteria heavily produce numerous hydrolytic and transport proteins targeting organic phosphorus components in response to phosphate limitation. Thus, exoproteomics can generate significant insights into the mechanisms utilised by microbes to compete for growth limiting nutrients and their contribution to environmental nutrient cyling^25, 29, 30^. In this study, we adapted our previous extraction method to efficiently capture the extracellular proteins (meta-exoproteome) found in agricultural soils to identify the most active microbial taxa in the rhizosphere of *Brassica napus* L. (Oil Seed Rape) and the major metabolic interactions operating. We hypothesised that 1) the rhizosphere would contain a distinct set of metabolically active microbes relative to the surrounding bulk soil and 2) microbes would express proteins for the mineralisation of N and P as a response to elevated C. In addition to capturing extracellular plant and aphid-pest proteins in the rhizosphere, we observed greater microbial activity in this compartment relative to the bulk soil with several *Pseudomonas* spp. dominating the metaexoproteome.

## Materials and methods

### Growth conditions for laboratory pot experiments

Plants were grown in a field soil collected from the University of Reading’s Sonning Farm facility (51° 28’ 55.3836” N, 0° 53’ 44.3688” W) between a depth of 20 and 60 cm. The soil properties are 50% sand, 38% silt and 12 % clay, with a pH of 6.7. Soil was air dried for 72 hr, sieved through a 1 cm sieve and supplemented with 0.4 g L^−1^ NH_4_NO_3_, 0.75 g L^−1^ KNO_3_ and 0.225 g L^−1^ Ca(H_2_PO_4_)_2_. Pots were filled with 1 L of soil and a 1 cm layer of moist perlite overlaid for the seeds to germinate in. Two seeds of *Brassica rapa* R-o-18 were sown in to the perlite layer to germinate, and thinned to one plant per pot. At sowing pots were treated with one of the following 1) 100 mL minimal media, identical to^27^; 2) 100 mL minimal media with *Pseudomonas putida* BIRD-1. *P. putida* BIRD-1 was maintained on Luria Bertani (LB) agar (1.5% w/v) medium at 30°C. Prior to inoculating the pots, BIRD-1 was grown overnight in minimal medium using glucose as the sole carbon source^27^ to 10^9^ cells mL^−1^. The experimental design consisted of two treatment levels (+/− BIRD-1) with each treatment combination consisting of six replicate plants.

Pots were placed in large seed trays and watered from below using deionised water throughout the experiment. Pots were placed in a controlled environment growth room (Weiss Technik UK Ltd., Units 37-38, Loughborough, UK), with a 16 h day length, 21 °C day, 18 °C night and 80% RH. Light was provided by a bank of fluorescent bulbs with a PPFD of 250 mol m^−2^ s^−1^. After 4 weeks growth, plants were harvested. Plants were gently extracted from the soil and excess soil shaken off the root system. Roots were cut and placed in 50 mL falcon tube containing 20 mL potassium sulphate buffer (PSB; 0.5M K_2_SO_4_, 10 mM EDTA, pH6.6). The roots were vortexed in the PSD for 10s to remove rhizosphere soil before being transferred to a fresh tube containing PSB and vortexed again for 10 s. The rhizosphere soil from both tubes was combined into one sample, centrifuged at 4000 rpm at 4 °C for 5 min and then snap frozen in liquid nitrogen. Samples were stored at −80 °C prior to being freeze dried.

### Field sampling site and conditions

*Brassica napus* L. plants were sampled at the four-leaf growth stage in October 2017 from the same location as the soil was collected for the pot trials above. Plants were removed from the soil and the roots processed as described above.

### Extraction of extracellular proteins from soil

To extract extracellular proteins from agricultural field soil, the methods developed by Johnson-Rollings et al (2014) were modified to account for the reduction in available sample associated with rhizosphere soil. Briefly, loose soil was shaken off plant roots and discarded, and the remaining rhizosphere soil was removed from the roots by immersion and shaking in a 0.5M KSO_4_ 10mM EDTA buffer, pH 6.6, until approximately 30g of soil had been collected, in a 1:3 w/v ratio of soil: buffer. This solution was incubated at room temperature with 100rpm shaking for 1 hour, and centrifuged at 12800xg for 20 min at 4°C, decanted into Nalgene centrifuge tubes and centrifuged at 75600xg for 20 min at 4°C. The supernatant was then sequentially filtered through 0.45 and 0.22μm pore-size PVDF filters (Fisher Scientific) to remove any bacterial cells and adjusted to pH 5 with 10% v/v Trifluoroacetic acid. 0.001% (v/v) of StrataClean resin (Agilent) was added in order to bind protein, and samples were incubated in a rotatory shaker at 4°C overnight. Samples were centrifuged at 972xg for 5 min at 4°C, and supernatants were discarded. If any buffer had crashed out of solution, then the resin was resuspended in dH2O adjusted to pH 5 with 10% v/v Trifluoroacetic acid, and this centrifuge step was repeated. Next, the resin was resuspended in 20μl of 1xLDS 1xDTT gel loading buffer (Expedeon), and heated to 95°C for 5 min, then sonicated in a water bath for 5 min, twice in succession.

### Identification and quantification of proteins

For protein identification a short run (~2 min) was performed to create a single gel band containing the entire exoproteome, as previously described by Christie-Oleza et al., (2012)^31^. In-gel reduction was performed prior to trypsin digestion and subsequent clean up as previously described by ^31^. Samples were analysed by means of nanoLC-ESI-MS/MS using an Ultimate 3000 LC system (Dionex-LC Packings) coupled to an Orbitrap Fusion mass spectrometer (Thermo Scientific, USA) using a 60 min LC separation on a 25 cm column and settings as previously described^32^.

To identify peptides, we used an iterative database search approach. First, all detected mass spectra were searched against the total assembled MG database containing 65 M open reading frames (ORF), generated from a composite metagenome of the field soil, detailed below. To reduce redundancy, ORFs were clustered at 90% using CD-HIT and representative ORFs sequences were used as the database. X!Tandem and MS-GF+ searches were performed, generating a database of 206065 identified proteins, prior to FDR and minimum unique peptide filtering. This reduced ORF database was then used in a MaxQuant search, returning 6718 proteins (plus 71 decoy and 21 contaminants). Removal of proteins with only one observed peptide, only identified by modified peptides, and allowing for a FDR of 10% resulted in a final protein detection of 1895 protein groups. The highest ranked protein in each group, based on number of unique peptides and/or probability was taken forward. Typically, protein groups consisted of proteins of identical function separated by taxa, predominantly at the species level. Quantification, statistical analyses and data visualisation of exoproteomes was carried out in Perseus^3^ and Rstudio (version 1.2.5033). The mass spectrometry proteomics data have been deposited in the ProteomeXchange Consortium via the PRoteomics IDEntifications (PRIDE) partner repository with the dataset identifier (TBC).

### Extraction of metataxonomic and metagenomic data

DNA from either bulk or rhizosphere soil was extracted using the FastDNA™ Spin Kit (MP Biomedicals™) soil extraction kit following the manufacturer’s instructions. All samples were checked for integrity and quality by gel electrophoresis (1% w/v agarose) and NanoDrop Spectrophotometry (ThermoFischer). DNA was quantified using QuBit (ThermoFischer). For 16S rRNA gene profiling of the microbial communities 16S rDNA amplicons covering the V1-3 variable regions were amplified using 27F and 534R eubacterial primers with Illumina overhang adapter sequences. Following PCR cleanup (as per the manufacturer’s instructions) using AMPure XP beads (Beckmann Coulter), indices were attached using the Nextera XT index kit (Illumina) as per the manufacturer’s instructions. Amplicons were quantified, pooled, and prepared for 2×300bp paired end sequencing using an Illumina Miseq platform, as per the manufacturer’s instructions. For shotgun-metagenomes, libraries and sequencing was performed by Novagene Ltd using an Illumina HiSeq – PE 150 bp.

Metataxonomic assessment of microbial communities using the 16S rRNA marker gene was performed using QIIME2 (Ver. 2020.11)^1^. Singleend (forward reads) files were demultiplexed using the demux plugin. Then, quality control was performed on each sample using the Dada2 plugin ^2^. All amplicon sequence variants (ASVs) were aligned with mafft and used to construct a phylogeny with fasttree2 (via q2-phylogeny) ^3^. Taxonomy was assigned to ASVs using the q2-feature classifier against the Greengenes 13_8 97% reference sequences^4^. Data from genome sequencing has been deposited in the NCBI Sequence Read Archive (SRA) under accession number XXXXXX.

A Bray-Curtis dissimilarity matrix was calculated based on each samples clustered-ASV profiles and used for non-metrical multidimensional (NMDS) scales. We have modelled the distances between UniFrac and the Bray – Curtis discrepancies using the ASV-Level table through one-way similitude analysis (ANOSIM) to investigate differences in community composition between rhizosphere and soil compartments. All of the above models were constructed using R-Studio (Version 3.6), including analyses for comparing the relative abundance of different bacterial taxonomic levels.

### Co-assembly of the composite metagenome

As a first exploratory step, different co-assemblies schemes of were explored, grouping samples by fertilizer or phosphate content categories as well as a full co-assemblies. From simple statistics such as N50 and total assembly size, the full co-assembly was chosen for downstream analysis. All assemblies were carried out using megahit [62] version 1.1.3. After removing contigs smaller than 500 nucleotides, open reading frames (ORFs) were called using prodigal version 2.6.2 [63] with option “-p meta”. This resulted in a collections of 64.1 millions different ORFs. ORFs multi-sample coverage profiles were generated by mappings all samples reads to assembly using both bwa-mem version 0.7.17-r1188 [64] and samtools version 1.10 [65].

### Metagenomic Taxonomic profiling

Assembly of the 16S rRNA gene does from shotgun metagenomic read data is notoriously poor and susceptible to high variable copy number across bacterial taxa. Therefore, community composition of the MG was based on the taxonomy and abundance of **S**ingle copy **C**ore **G**enes (SCG). For this, the database of 64 millions ORFs was annotated using rpsblast version 2.9.0+ [66] using the pssm formatted COG database [67], which is made available by the CDD [68]. For the set of 36 COG taken as SCG, corresponding ORFs were clustered at 5% ANI using mmseqs2 version 13.45111 [69] with options “easy-cluster”, “--cov-mode 2”, “-- max-seqs 1000” and “-c 0.80”. This resulted in a median number of 6280 clusters over the 36 SCGs. After adding sequences from Refseq genomes representatives, a series of 36 corresponding phylogenetics trees were built using in sequence, mafft version v7.407 [70], trimal version v1.4.rev22 [71] with options “-gt 0.9” and “-cons 60”, and FastTree version 2.1.10 [72]. Using the python library ete3 version 3.1.2 [73], each SCG cluster were assigned to the nearest refseq representative. Taxonomic profiles were obtained by summing SCG cluster’s ORFs coverages along taxonomic assignment. In the case where more than one of the 36 SCG was found at the same taxonomic level, median coverage was taken. Normalisation was carried out by taking for each sample the median total coverage of the 36 SCG. This approach is insensitive to the varying size of genomes, so that organisms with larger genomes do not appear to be more abundant than those with smaller genomes.

To identify individual phosphatases (PhoX, PhoD and PhoA) the methods developed in ^33^ were applied. Briefly, this involved performing a hmmsearch of the assembled MG ORF database, using generated profile hidden markov models for each protein. To assign taxonomy, identified ORFs were then aligned by BLASTP (e^−20^) to a manually curated database generated from all sequences deposited in the IMG/JGI database. Any sequences (<2%) not aligning to any predicted phosphatases were removed from the dataset. The read coverage for each ORF was then extracted to determine the relative abundance of each ORF. ORF coverage was then normalised by accounting for variation in total SCG coverage as per Murphy et al. (2021). For PhoX, phylogenetic analyses were performed using IQ-Tree using the parameters -m TEST -bb 1000 -alrt 1000. Representative PhoX sequences obtained from the genomes of isolates spanning the diversity of *Pseudomonas* were aligned with the sequences identified in the meta-exoproteome. Evolutionary distances were inferred using maximum-likelihood analysis. Relationships were visualised using the online platform the Interactive Tree of Life viewer (https://itol.embl.de/).

## Results

### *In situ* metaexoproteomics reveals *Pseudomonas putida* BIRD-1 expresses a distinct set of rhizosphere-associated proteins under laboratory conditions

Our previous extraction method involved extracting protein from ~100g soil^26^, which is not feasible when working with rhizosphere soil. To determine whether we could efficiently capture extracellular proteins from soil using the Stratabead resin we first performed a series of protein spikes into soil and water as a control. We used either bovine serum albumin or the exoproteome of lab-grown *P. putida* BIRD-1 as the protein spike, both of which were successfully re-captured from soil and water (Fig. S1). To further confirm the efficacy of this method, we established a simple plant growth experiment under laboratory conditions using *Brassica rapa* OH17 grown in a sand: soil mix (n=6) and inoculating washed and resuspended *Pseudomonas putida* BIRD-1 (6×10^8^ cells ml^−1^). After three weeks growth, the sand: soil mix was collected, and protein extracted.

Using the *in silico* predicted proteome of either *P. putida* BIRD-1 or *B. napus* as the database and a criterion of at least 2 unique peptides per protein, we identified a total 201 (177 protein clusters) and 215 proteins, respectively (Table S1 & S2). Two replicate samples containing low quantities of protein were omitted from further analyses. For *P. putida* BIRD-1, the 40 most abundant proteins represented 61% of the total exoproteome. The pot-grown *P. putida* exoproteome (Black bar) showed a distinct profile compared to previous *in-vitro* exoproteomes^27^ (Blue bar, Figure 1A, Table S1)). This included an increase in the relative abundance of some outer membrane proteins, porins and substrate binding proteins associated with ABC transport systems and a decrease in others, whilst some were detected at similar relative abundances (Table 1). Whilst we did detect several cytoplasmic proteins, their relative abundance in either the exo- or intracellular soluble-(Grey bar) proteomes were still markedly different. In addition, many of the abundant *in-vitro* intracellular proteins detected in our previous study^27^ were either not detected or were present in very low abundance in the pot-grown proteome suggesting an inherent change in metabolism. Most of the abundant rhizosphere-inducible proteins belonged to four general COG categories: 1) Amino acid transport and metabolism 2) Carbohydrate transport and metabolism 3) Inorganic ion transport and metabolism 4) Cell envelope biogenesis, outer membrane (Fig. 1B, Table 1). Whilst several substrate binding proteins related to carbon and nitrogen metabolism were enriched in the rhizosphere (Fig. 1C), substrate binding proteins associated with high-affinity phosphate (PstS) or 2-aminophosphonate (AepX) ABC transporters were not detected, nor was the low-phosphate inducible alkaline phosphatase PhoX^27^. This would suggest that under these conditions BIRD-1 did not experience localised phosphate depletion severe enough to trigger its P-stress response regulon^27^.

**Table 1.**
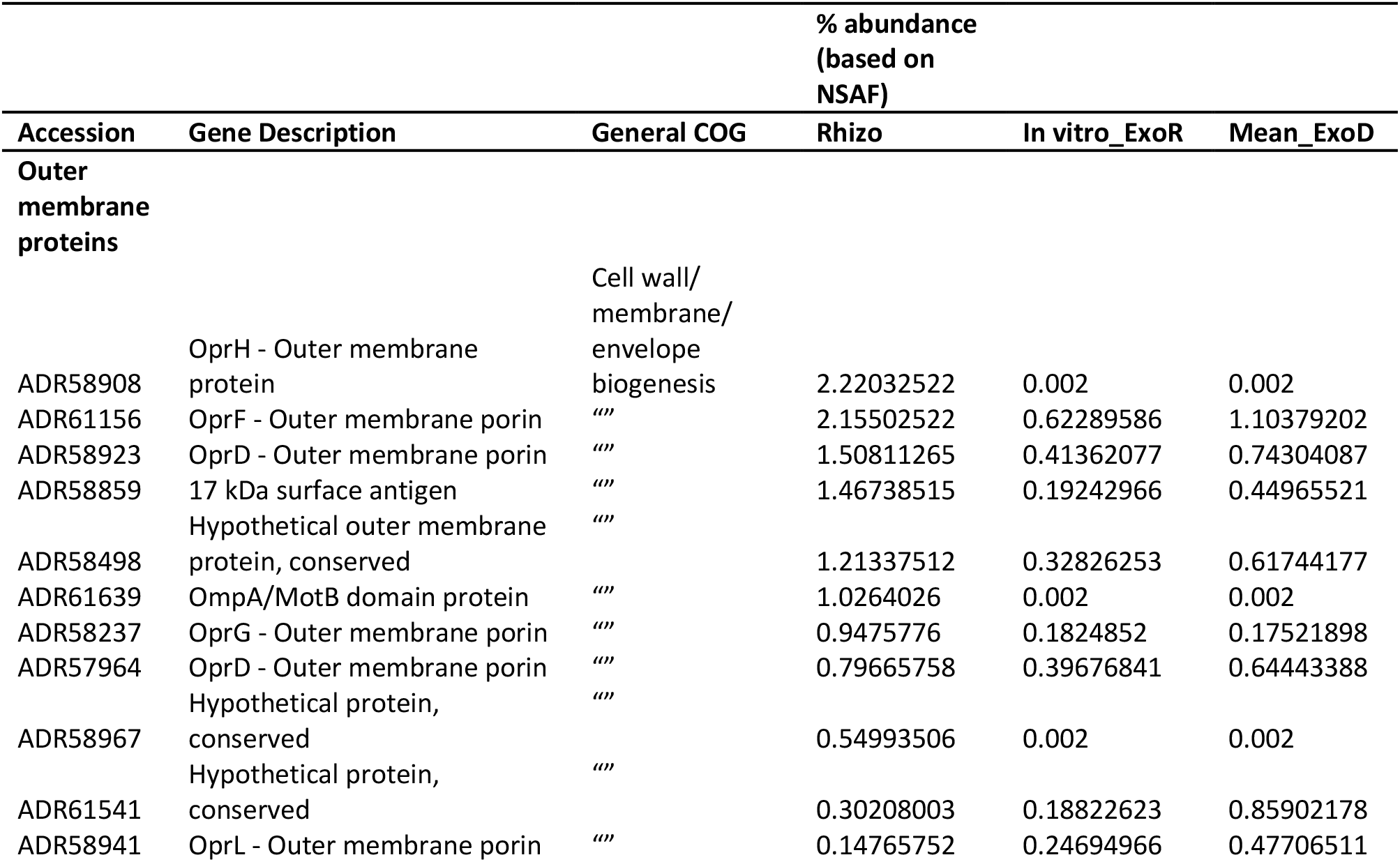
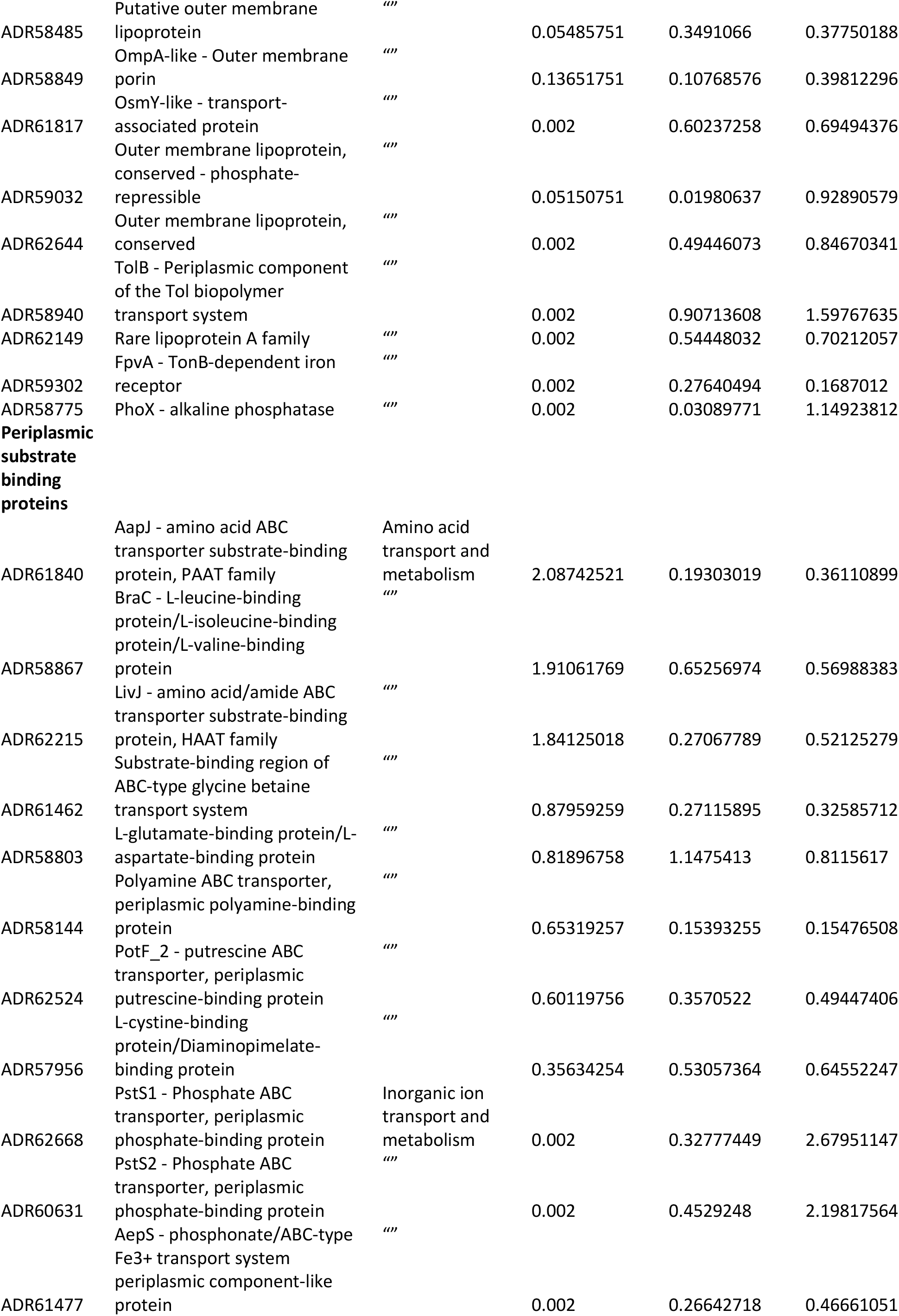
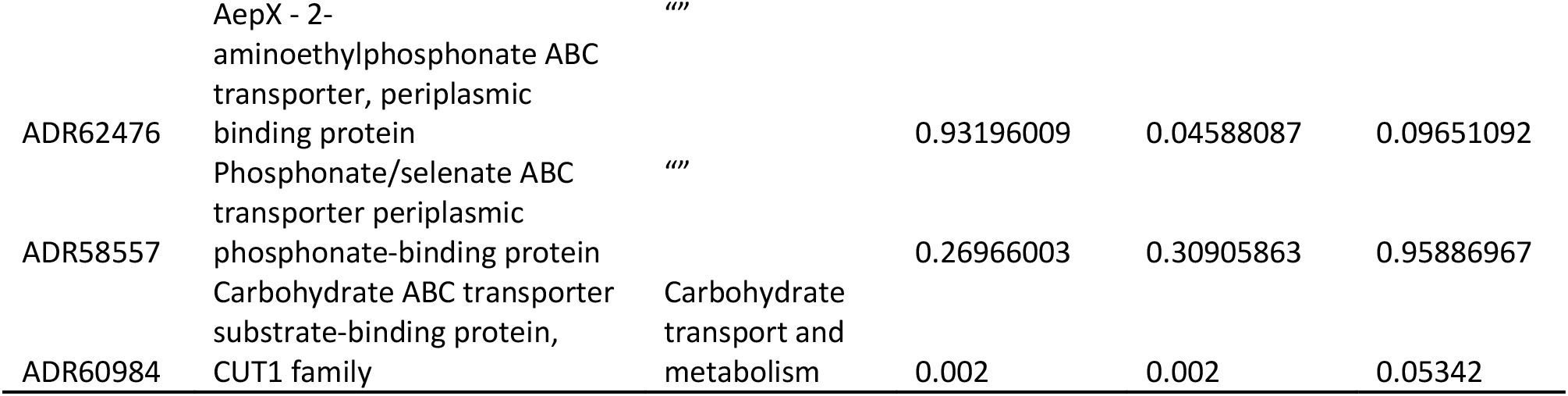
Comparison of extracellular proteins with validated/predicted outer membrane or periplasmic localisation detected in the exoproteome of *P. putida* BIRD-1 grown in *B. rapa* 0H18 rhizosphere soil (Mean_pot), *in-vitro* (phosphate replete (Mean_ExoR) and phosphate deplete (ExoD) growth conditions). Values present are the calculated mean % abundance (n=4) in the total exoproteome based on normalised spectral abundance factor (NSAF) values.

**Figure 1.**
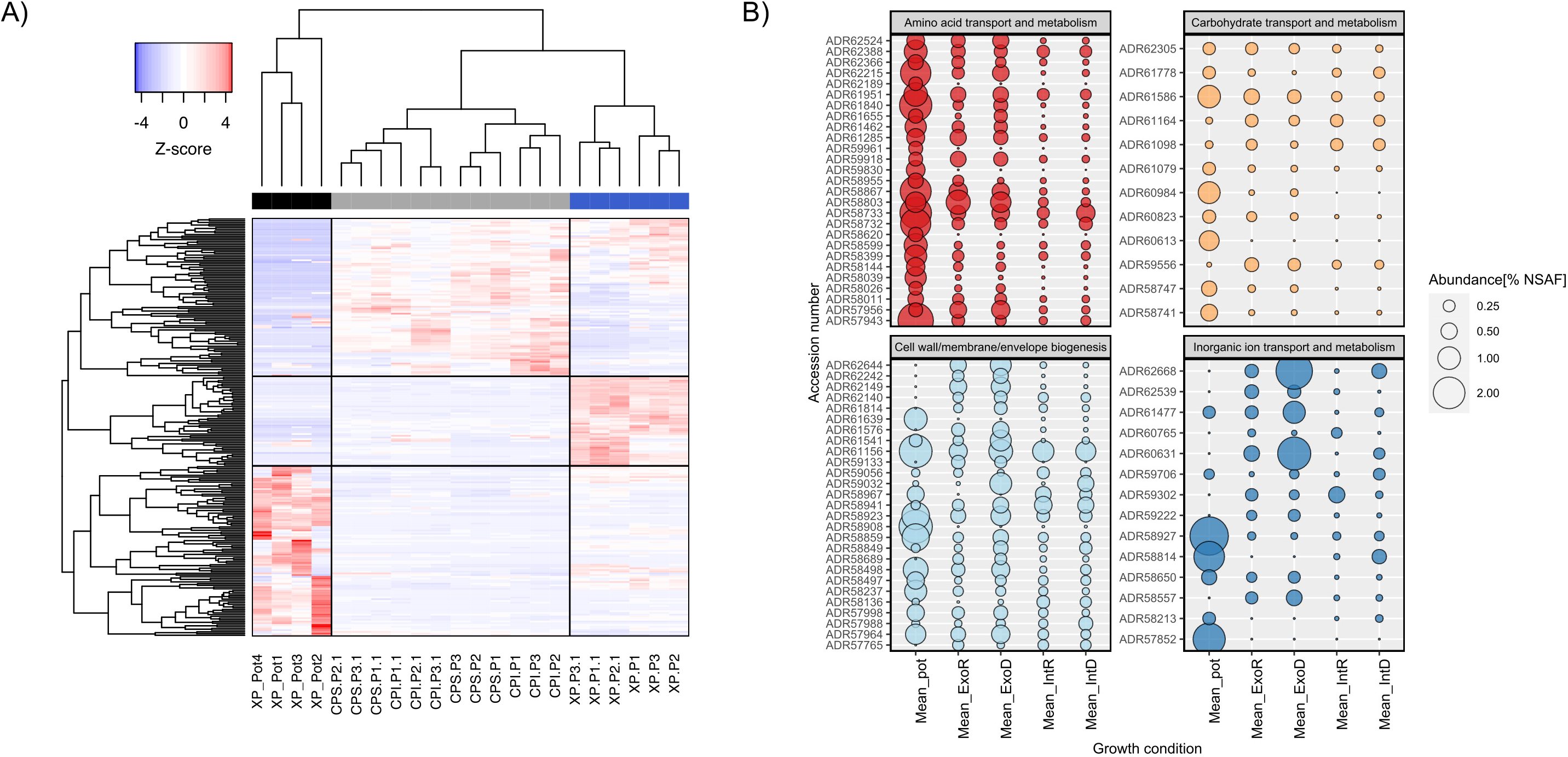
Comparison of *P. putida* BIRD proteome during *in situ* and *in vitro* growth experiments. **(A)** Hierarchical clustering of proteins based on abundance profiles (Z-score, calculated from % abundance) across the different growth conditions; blue, exoproteome (XP) of liquid cultures; Grey, insoluble and soluble fractions of the cellular proteome (CP) from liquid cultures; black, exoproteome captured from pot experiments using *Brassica rapa* R018. Data for individual replicates is displayed **(B)** Assessment of functions (COG categories) related to periplasmic, cell surface and extracellular proteins across the different growth conditions. The mean value for quadruplicate (pot), triplicate (In vitro deplete/replete) and sextuple (In vitro deplete/replate) replicates from each growth conditions is plotted. GenBANK accession number are given on the y axis. All % abundance values were calculated from Normalised Spectral Abundance Factor (NSAF) values.

### Metaproteomic assessment of microbial activity in the rhizosphere of field-grown Oil Seed Rape and surrounding bulk soil

Next, we sampled bulk and rhizosphere soil from agricultural fields sown with Oil Seed Rape under contrasting P fertiliser regimes (Fig. S2). Plants were sampled at an early growth stage, between the four-six leaf and rosette growth stages. To create a comprehensive database for meta-exoproteomics, we generated over 500 GB of metagenomic data from bulk and rhizosphere soil samples and used a co-assembly method to identify over 64 million open reading frames (ORFs). When controlling for one of the two variables separately, nonmetric multidimensional scaling (NMDS) analysis showed no significant effect of compartments (bulk v rhizosphere; ANOSIM: R=0.027, p = 0.316) or fertiliser treatment (High P v Low P; ANOSIM: R=0.4852, p = 0.11) on the total soil microbial communities (MG) (Fig. 2A & B). Again, using a minimum of two unique peptides per protein, a total of 1881 proteins were detected across all samples and there was a significant difference (Adonis: R^2^=0.736, P = 0.001) in the proteomic profiles of bulk and rhizosphere samples (Fig. 2C). In contrast, there was no significant effect (Adonis: R2=0.031, P = 0.667) of fertiliser treatment on the proteomic profiles of either soil compartment. Whilst only 42 proteins were significantly enriched in the bulk soil (FDR corrected P<0.05), over 500 were significantly enriched (FDR corrected P<0.05) in the rhizosphere (Table S3). Furthermore, almost all highly abundant proteins were rhizosphere associated, with only few abundant proteins associated with bulk soil (Fig. 2D).

**Figure 2.**
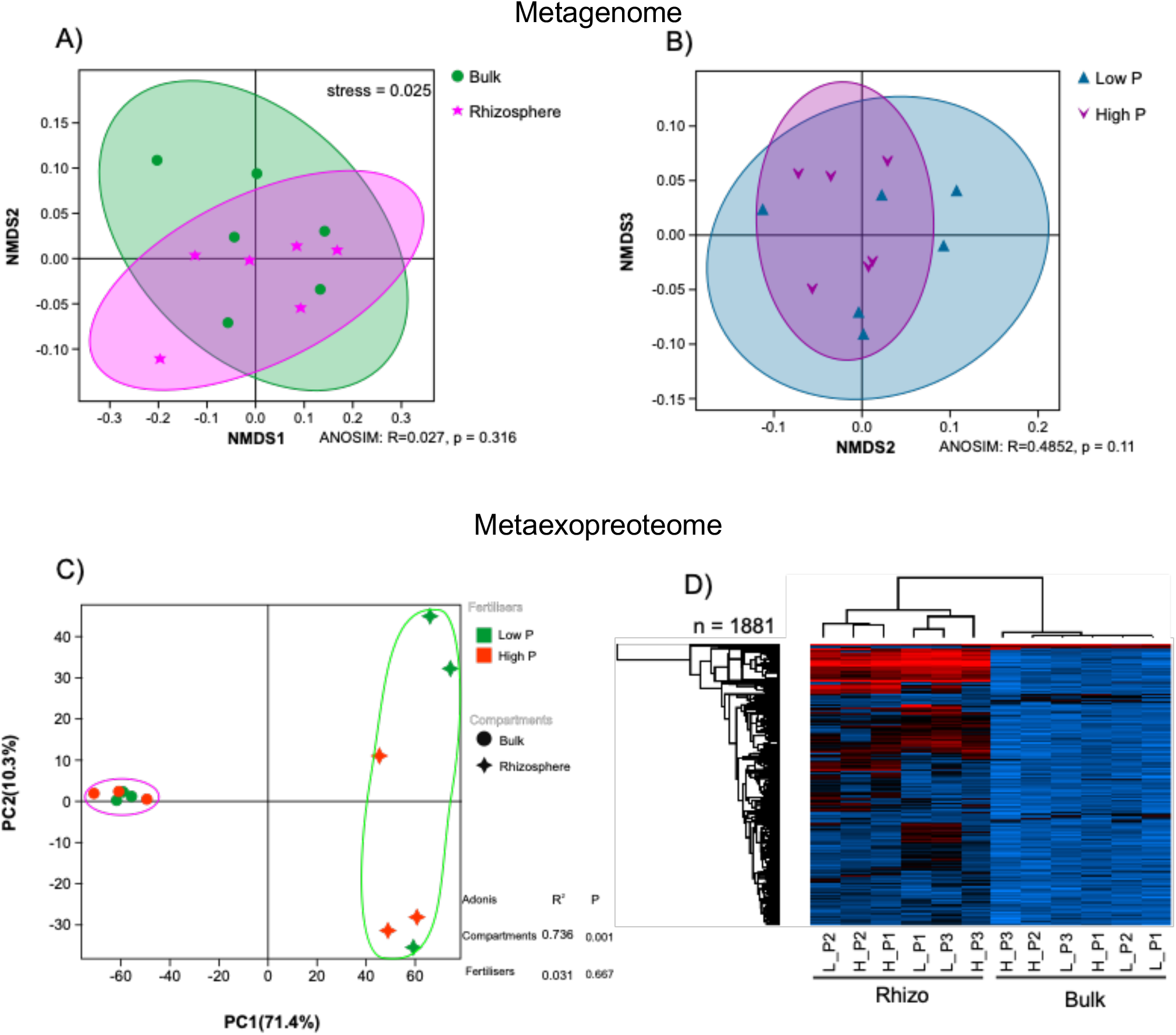
Metagenomic and metaexoproteomic assessment of field-grown *Brassica napus* L. bulk soil and rhizosphere communities. NMDS ordination (stress = 0.025) between total microbial community composition of **(A)** the rhizosphere (n=6) and bulk (n=6) compartments or (B) the low Pi fertilizer (Low P) and high Pi fertilizer (High P) treatments. Each point represents one metagenomic sample (n = 12). Data representing relative variable importance (R) and significance (p) calculated by PERMANOVA (Anosim) are displayed. **(C)** Multivariate analysis of the active microbial communities collected from the same soil samples, bulk soil (circles) and rhizosphere soil (squares). **(B)** The relative abundance of detected proteins in all samples based on label-free quantification (LFQ) values. Pale blue equals the least abundant, black equals the mean abundance and red equals the most abundant. Dendrograms for both sample and protein were calculated.

### Autochthonous *Pseudomonas* spp. were highly active in the rhizosphere of young field-grown *Brassica napus* L

The vast majority of rhizosphere protein content was related to several *Pseudomonas* spp. (~65%) and *B. napus* (~20%) (Fig. 3A). In addition, several abundant rhizosphere proteins were related to soil/root aphids and its corresponding symbiont *Buchnera aphicola* (*Gammaproteobacteria*). Proteins expressed by these groups as well as *Betaproteobacteria*, other *Gammaproteobacteria*, *Bacteroidetes* (predominantly *Flavobacteraceae*) and fungi were all rhizosphere-enriched (Fig. 3A). In contrast proteins expressed by *Actinobacteria*, *Alphaprotebacteria*, *Acidobacteria* and *Archaea* were collectively more abundant in bulk soil (Fig. 3A). The identified *Pseudomonas* proteins were aligned to four different *Pseudomonas* genomes, *P. putida* BIRD-1, *P. fluorescens* SBW25, *P. stutzeri* DSM4166, and *P. syringae* DC3000. On average, detected proteins had the highest identity with *P. fluorescens* SBW25 (93%). However, there was significant variation in average identity (%) related to each strain and numerous proteins were absent from each individual genome (Fig. S3, Table S4), suggesting multiple strains of *Pseudomonas* were highly active in the rhizosphere.

**Figure 3.**
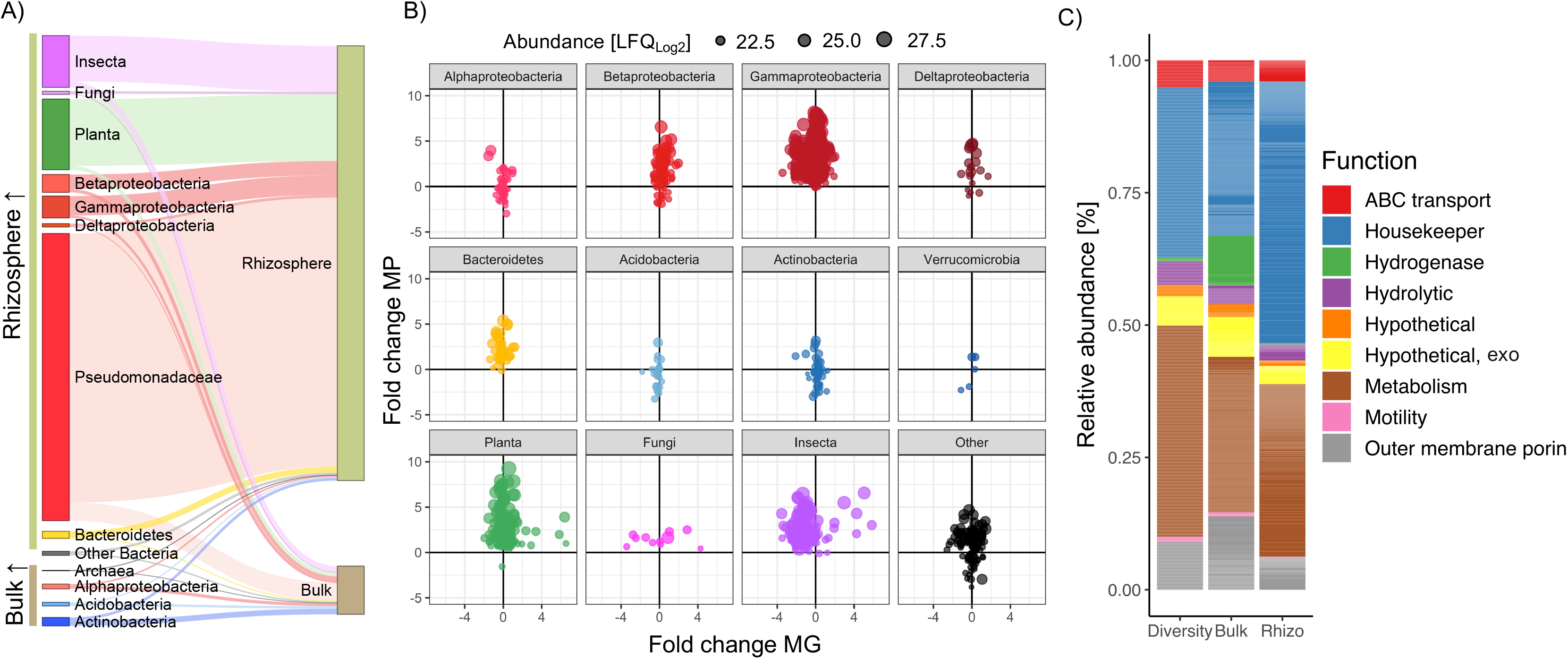
Taxonomic profile of the *in situ* meta-exoproteome sampled from field-grown *Brassica napus*. **(A)** Compartmental partitioning based on the relative abundance of proteins associated with various taxonomic groups identified in either rhizosphere or bulk soil samples. **(B)** Comparison of relative protein expression versus relative gene abundance in rhizosphere versus bulk soil samples. Each point represents a single protein and its size represents its relative abundance in the meta-exoproteome. Proteins were partitioned into various taxonomic groups. **(C)** Broad functional assessment of the bacterial meta-exoproteome. The relative abundance of these functions was calculated by either counting the total number of distinct detected proteins associated with each function (Diversity) or by determining their relative abundance (LFQ values) in either the bulk (Bulk) or rhizosphere (Rhizo) meta-exoproteome. Results plotted are the mean of 6 replicates: 3 Pi-replete and 3 Pi-deplete for each compartment.

Comparison of individual protein abundance with its corresponding ORF abundance in the MG demonstrated plant-associated bacteria, such as *Gammaproteobacteria* (*Pseudomonadaceae*), *Betaproteobacteria* (*Burkholderia, Oxalobacteraeae*, *Commonamondaeae*) and *Bacteroidetes* (*Flavobacteriaeae*) were more active in the rhizosphere despite minor changes in gene relative abundance (Fig. 3B). In addition, proteins expressed by methylotrophic bacteria were also enriched in the rhizosphere relative to surrounding bulk soil. For *Acidobacteria* and *Actinobacteria*, whilst some proteins were more abundant in the rhizosphere, other proteins were more abundant in the bulk soil. This is consistent with the life history strategy of many taxa related to these two phyla^34^. Proteins related to various genera associated with the Candidate Radiation Phyla were also detected demonstrating this large group of enigmatic bacteria are active in natural soils. Together, this demonstrates meta-exoproteomics can identify changes in metabolic activity that can be masked by solely relying on metagenomic data.

Based on functional annotation of all identified bacterial proteins, a large proportion of intracellular proteins were still captured during our extraction step, many of which were related to either central or auxiliary metabolism as well as housekeeping functions and protein synthesis (Metabolism, Housekeeper, Fig. 3C). Despite the greater amount of total protein detected in rhizosphere samples (Fig. 2D), ribosomal proteins (protein synthesis marker) represented a greater proportion of total protein in this compartment, further demonstrating elevated microbial activity in this rhizosphere compartment relative to bulk soil. The most abundant extracellular proteins were related to outer membrane porins, substrate binding proteins associated with ABC transporters and extracellular hydrolytic enzymes. In bulk soil, but not in the rhizosphere, a significant proportion of protein was assigned to hydrogenases.

### Taxonomic assessment of total soil communities associated with field-grown *Brassica napus* L

Microbial community composition did not differ significantly between compartment (Fig. 2A) and many rhizosphere-enriched proteins were encoded from genes whose abundance in the MG showed little variation between either soil compartment (Fig. 3B). Therefore, to better determine the relative abundance of active taxa identified in the MP we expanded our taxonomic assessment of the total soil microbial communities (MG) associated with either bulk soil or rhizosphere soil compartments. To do this, we analysed the assembled MG, using the read abundance and taxonomy of SCGs, as well as generating an amplicon-based 16S rRNA gene profile (Fig. 4). Both the SCG and 16S rRNA gene profiles showed Actinobacteria (MG = 24%; 16S bulk = 48%, 16S rhizo = 43%) and *Proteobacteria* (MG = 44%; 16S bulk = 21%, 16S rhizo = 29%) numerically dominated the total genomic content of the total soil bacterial community, whilst *Bacteroidetes* constituted only 5%. At the order-level, based on SCGs, separation by compartment (bulk v rhizosphere) did not significantly affect the soil communities’ taxonomic structure (Fig. S4). 16S rRNA gene profiles revealed the abundance of *Pseudomonadaceae* (3.426%; 100% *Pseudomonas*) was not significantly different between bulk and rhizosphere compartments, despite significantly greater activity in the rhizosphere, as was the case for SCG-derived abundance data (Fig. 3A). In contrast, rhizosphere-active *Flavobacteraceae* (100% *Flavobacterium*) and *Betaproteobacteria*, which made up 3.804% and 6.12% of the 16S rRNA gene rhizosphere community, respectively, were significantly enriched in this compartment compared to the bulk soil (Fig. S5). In summary, two methods for taxonomic assignment and relative abundance determination demonstrated significantly less shifts between soil compartments based on DNA in comparison with protein.

**Figure 4.**
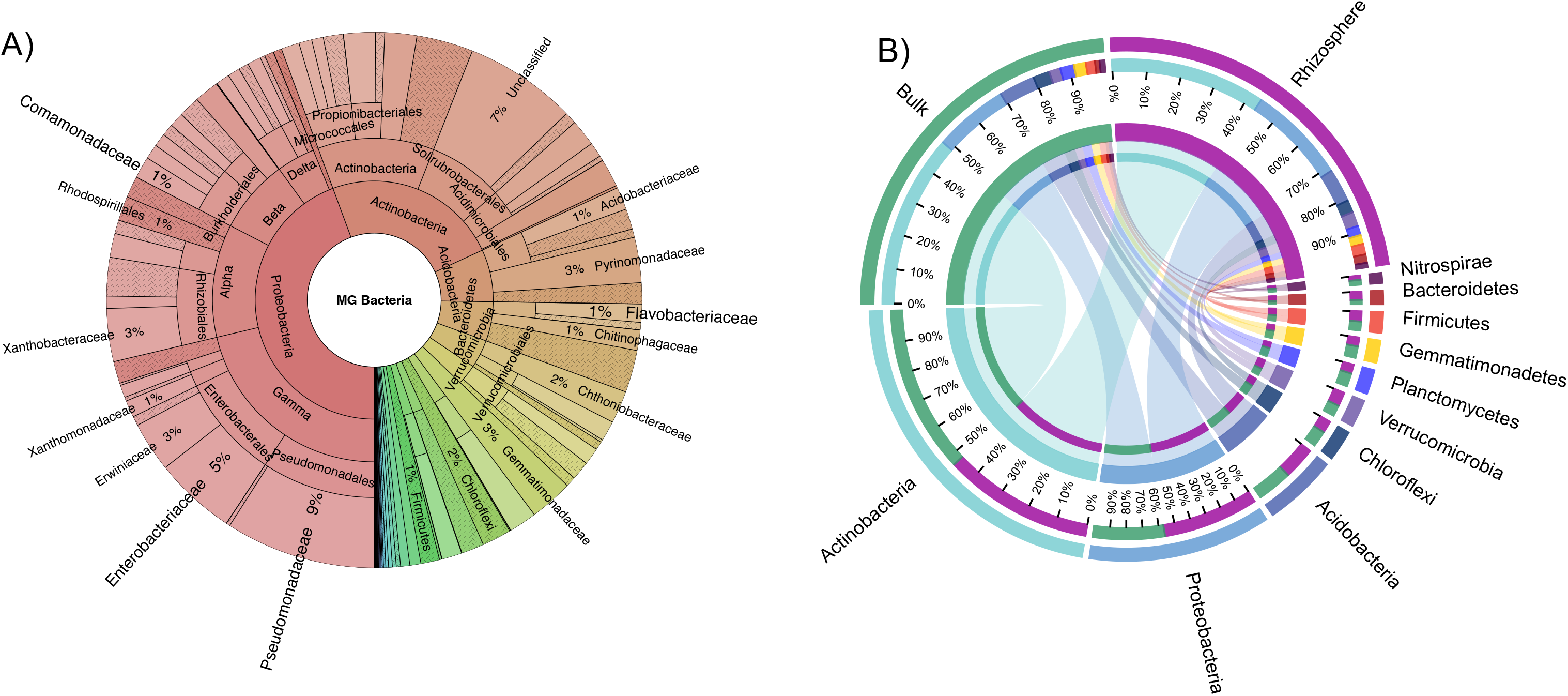
Composition of bulk soil and rhizosphere soil microbial communities sampled from field-grown oil seed rape based on the composition of single copy core genes in the metagenome or 16S rRNA gene amplicon profiling. **(A)** Relative abundance of all bacterial taxa in the combined (bulk and rhizosphere) soil metagenome. Selected taxonomic groups of interest in this study are labelled whilst others have been omitted for clarity. **(B)** CIRCOS plots showing the relative abundance distribution among the dominant phyla in either the bulk soil or rhizosphere compartment.

### Active autochthonous *Pseudomonas* spp. experience Pi-limitation under field conditions independently of fertiliser regime

To better determine the metabolic interactions occurring in the rhizosphere, we focused our analysis on the proportion of proteins predicted to be either periplasmic, outer membrane associated, or extracellular, ignoring predicted cytoplasmic proteins whether they had a role in nutrient acquisition of not. The expression of ABC-transporter related substrate binding proteins is an excellent proxy for metabolic interactions operating in any given environmental niche. The majority of these were expressed by *Pseudomonas* spp., and to a lesser extent *Burkholderiales* spp. (Fig. 5A). In agreement with our laboratory pot experiment inoculated with *P. putida* BIRD-1, many detected substrate binding proteins were associated with predicted amino acid transporters, but a significant number were also predicted to transport other nitrogenous compounds such as polyamines and quaternary amines, as well as carbohydrates (Fig. 5A, lower panel). Importantly, in the rhizosphere compartment, 2/3 *Pseudomonas* proteins identified as PstS, the substrate binding protein associated with the high affinity phosphate ABC transporter, were highly expressed (Fig. 5A, 5B). Indeed, one identified PstS was among the top 10 most abundant proteins in the total MP, including abundant plant and insect proteins (Table S3). Whilst numerous substrate binding proteins were significantly more abundant in the rhizosphere, fertiliser regime had no significant effect on their expression, even for PstS (Fig. 5A, Table S3).

**Figure 5.**
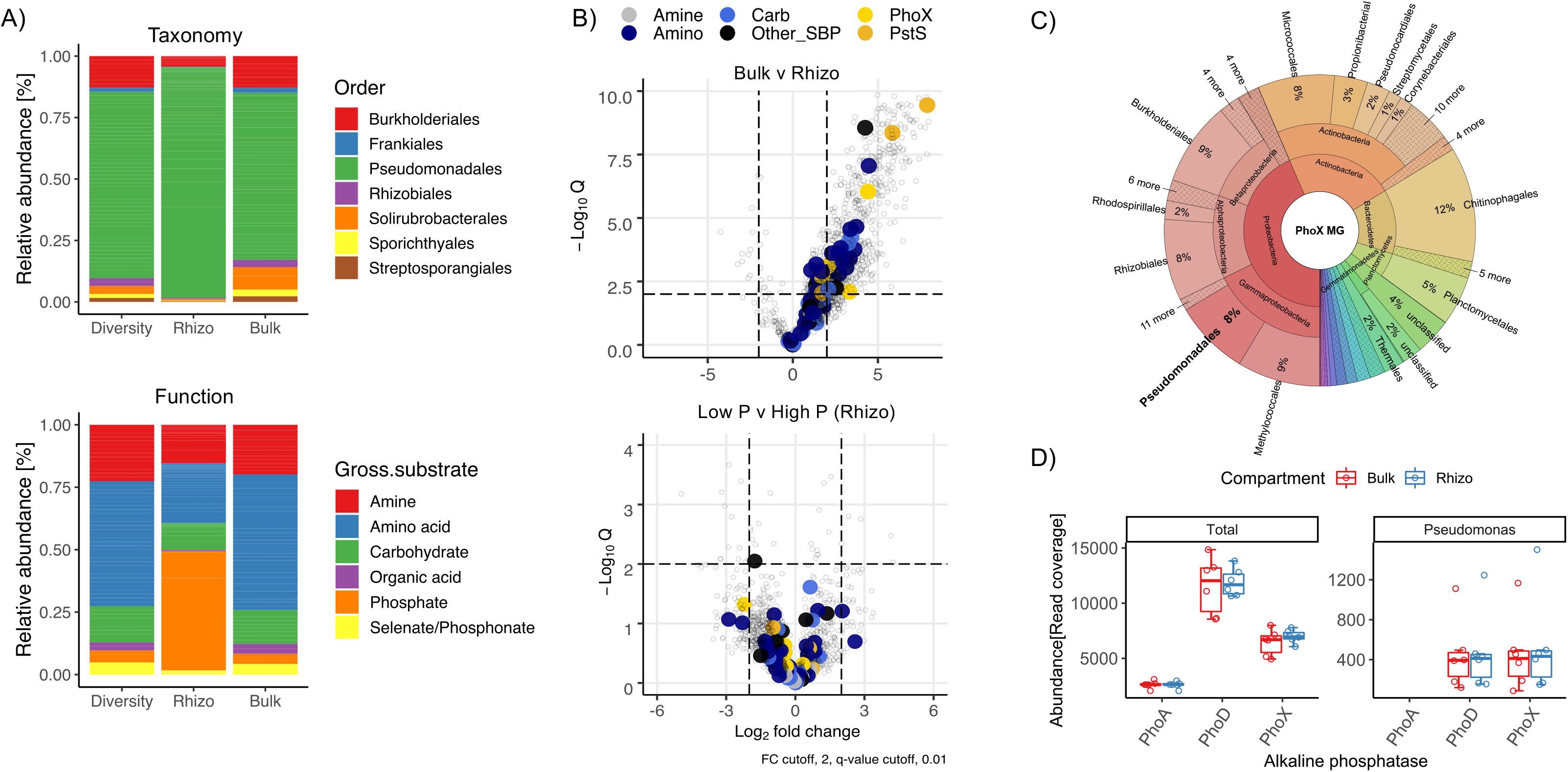
Functional analysis of *in situ* plant: microbe interactions based on soil metaproteomes. Taxonomy and gross functional classification of substrate binding proteins identified in bulk and rhizosphere (rhizo) soil metaproteomes. The relative abundance of each function was calculated by either counting the total number of distinct detected proteins associated with each function (Diversity) or by determining their relative abundance (LFQ values) in either the bulk (Bulk) or rhizosphere (Rhizo) meta-exoproteome. **(A)**. The effect of compartment and phosphate fertiliser regime on the expression of bacterial substrate binding proteins **(B)**. Taxonomy **(C)** and relative abundance **(D)** of all PhoX ORFs identified in bulk and rhizosphere soil MGs combined.

In addition, we also detected numerous extracellular hydrolytic enzymes (Fig. 5, Table S3). Among these were five alkaline phosphatases belonging to the PhoX family, all of which were most closely related to the *P. fluorescens* group (Fig. S6). Despite the limited diversity of alkaline phosphatases in the MP, *Pseudomonas* PhoX ORFs constituted 2% and 8% of the diversity and richness (relative abundance) in the MG, respectively (Figure 5C). Furthermore, despite their absence in the MP, there were also a significant number of ORFs related to either the PhoD or PhoA alkaline phosphatases in the MG, with PhoD the most abundant of all three (Fig. 5D). PhoA and PhoD homologs related to a diverse range of taxa were present in both compartments, (Fig. S6) despite no evidence of expression. The abundance of PhoD related to *Pseudomonas* was almost identical to PhoX related to *Pseudomonas* (Figure 5D), further suggesting preferential expression of the latter alkaline phosphatase.

## Discussion

Building a holistic understanding of plant:microbe interactions relies on the development of suitable tools to investigate complex and simultaneously occurring metabolic processes *in situ*. Here, first using *P. putida* BIRD-1 as a model and then analysing natural field-soil microbial communities, we demonstrate that meta-exoproteomic assessment of the rhizosphere is achievable and can significantly refine our understanding of the establishment and function of the plant microbiome. Specifically, these data generate further testable hypotheses surrounding the genomic basis of rhizosphere competence and microbial nutrient cycling, which will ultimately guide our ability to engineer plant microbiomes and better determine abiotic and biotic causes of plant disease. Likewise, this method successfully captured extracellular plant and corresponding pathogenic aphid proteins demonstrating the efficacy of this method to also understand plant host: pathogen interactions.

Plants differ in their ability to manipulate soil communities. For example, whilst a strong ‘rhizosphere effect’, i.e. enrichment of rhizosphere-specific bacteria recruited from the surrounding bulk soil, can be observed for Barley^16^, other plants elicit much more subtle differences^15^. A significant limitation of metagenomics is its inability to clearly identify the most active microbes and metabolic processes occurring in a specific environment, which is further compromised with the inclusion of subtle spatiotemporal parameters. Whilst our metagenomic data was consistent with the observation that Oilseed Rape elicits a weak ‘rhizosphere effect’ on soil microbial communities^14^, by utilising meta-exoproteomics we observed a clear difference between the active microbial community present in the rhizosphere compared to the surrounding bulk soil. This agrees with meta-transcriptomic studies investigating the active plant microbiota in various crops^35^. This difference in activity can be largely attributed to an increase in the quantity of microbial and plant protein captured in the rhizosphere, and elegantly demonstrates the rhizosphere as a hotspot for plant-induced microbial activity^1, 2, 36–38^.

Partitioning the meta-exoproteome between compartments was not only achieved through the capture of significantly more bacterial protein in the rhizosphere (Fig. 2B), but also a shift in the taxonomic groups producing these proteins. Rhizosphere-specialised bacterial taxa can also be thought of as copiotrophs, responding to elevated labile and complex organic carbon deposition. On the other hand, oligotrophs are relatively more active in carbon-depleted bulk soil^34^. The large increase in *Pseudomonas* activity as well other bacterial groups such as *Flavobacterium* (*Bacteroidetes*), and various *Burkholderiales* (*Betaproteobacteria*) in the rhizosphere is consistent with their predicted life history strategies^34, 39^. Likewise, our meta-exoproteomics data also confirmed that ‘bulk soil-specialised’ or oligotrophic bacteria, such as those related to *Verrucomicrobia*, *Actinobacteria* and *Acidobacteria^39^* were relatively more active in the surrounding soil. Whilst our study only captured a single time-point, our data clearly revealed that various distinct strains of *Pseudomonas* are highly active in the rhizosphere of Oil Seed Rape and represent a major and ecologically important component of this crop’s microbiome^14, 40, 41^. *Pseudomonas* represents a relatively small fraction of the seed microbiome, thus an increase in their relative abundance, especially several strains, in both the rhizosphere and root during early stage of OSR growth indicates active selection from the surrounding bulk soil^40, 41^.

In addition to gaining functional insight into the plant microbiome through identification of active taxonomic groups, we also identified numerous proteins related to various beneficial functions^1, 36^. Plant root exudates shape microbial communities and amino acids can become the major group of exudates released by plants over time and, in addition to quaternary amines, can represent a significant fraction of the dissolved organic N pool^42–44^. Furthermore, the turnover of the soluble amino acid pool in soil may be orders of magnitude greater than that of ammonium or nitrate^45^. Based on our MP data collected from both our inoculated pot experiment and field-trial we discovered *Pseudomonas* metabolism shifts towards the turnover of amino acids and other N-containing compounds when growing in the Oil Seed Rape rhizosphere. Given that most heterotrophic soil microbes are carbon-limited, the high expression of uptake and catabolic proteins targeting amino acids and other nitrogenous carbon sources (predominantly amines) observed here, suggests that microbial-mediated mineralisation of ammonium may be a key process in the rhizosphere, as observed in marine systems^30, 46^. Thus, release of nitrogenous organic carbon exudates may represent a mechanism that allows plants to get an immediate return on their metabolic investment in the form of labile ammonium. This aligns with the idea that plants ‘prime soils’ for microbial N mineralisation through the exudation of organic C, stimulating expression of peptidases and proteases^45^.

Plant-available phosphate is often a small fraction of the total soil P content. The slow diffusion of Pi in soil means that plant uptake during growth creates a zone of Pi-depletion around the roots (1-3mm)^47–50^, which is only intensified by increases in microbial growth on plant-derived labile organic carbon^1^. In almost all bacteria, including *Pseudomonas*, synthesis of phosphatases and the high affinity phosphate transporter PstSABC is negatively regulated by exogenous levels of inorganic orthophosphate. Thus, these proteins serve as excellent markers to assay for phosphate depletion^27, 51–53^. Furthermore, elevated soil phosphatase activity has recently been shown to co-occur with the severity of Pi depletion^54^. Whilst our pot experiments showed no evidence of localised Pi-depletion in the rhizosphere, in our field experiment the identification of five and three distinct *Pseudomonas* PhoX and PstS homologs suggests rhizosphere-dwelling *Pseudomonas* spp. experience phosphate-limiting growth conditions, despite saturation of the soil with inorganic fertilisers. PhoD is commonly used as the major gene marker for microbial phosphatase activity^55–57^. However, despite this family being the most abundant phosphatase in the MG for both the total community and *Pseudomonas* population, only PhoX was detected in the meta-exoproteome, consistent with its role as the major phosphatase in plant-associated *Pseudomonas^27, 53, 58^* and other environmental *Proteobacteria^59, 60^.*

## Conclusions

Here, we present the first meta-exoproteomic assessment of the plant microbiome sampled from a field-grown agricultural crop. Our new technique enabled us to identify highly active taxa in the rhizosphere and the key nutrients they target. Crop production heavily relies on the unsustainable use of inorganic N and P fertilisers and modern agricultural initiatives are moving towards the use of more sustainable organic sources of either N or P^61^. The success of this strategy is dependent on having a deep understanding of the key microbial players involved in N and P cycling and the biotic and abiotic factors which control this. In this regard meta-exoproteomics can greatly advance our understanding of the spatiotemporal dynamics of functionally important taxa and allow us to better engineer the plant microbiome through environmental and plant genotypic selection.

## Acknowledgments

We thank the Warwick Proteomics Research Facility, namely Dr. Cleidiane Zampronio for her assistance in generating and processing the mass-spectrometry data. This study was funded by the Biotechnology and Biological Sciences Research Council (BBSRC) and National Environmental Research Council (NERC) under project codes BB/L026074/1, BB/T009152/1 and NE/S013539/1 linked to The Soil and Rhizosphere Interactions for Sustainable Agri-ecosystems (SARISA) programme and a Discovery Fellowship (IL) and NERC Environmental ‘Omics Synthesis Grant (IL and EW), respectively.

---

**Figure S1.**
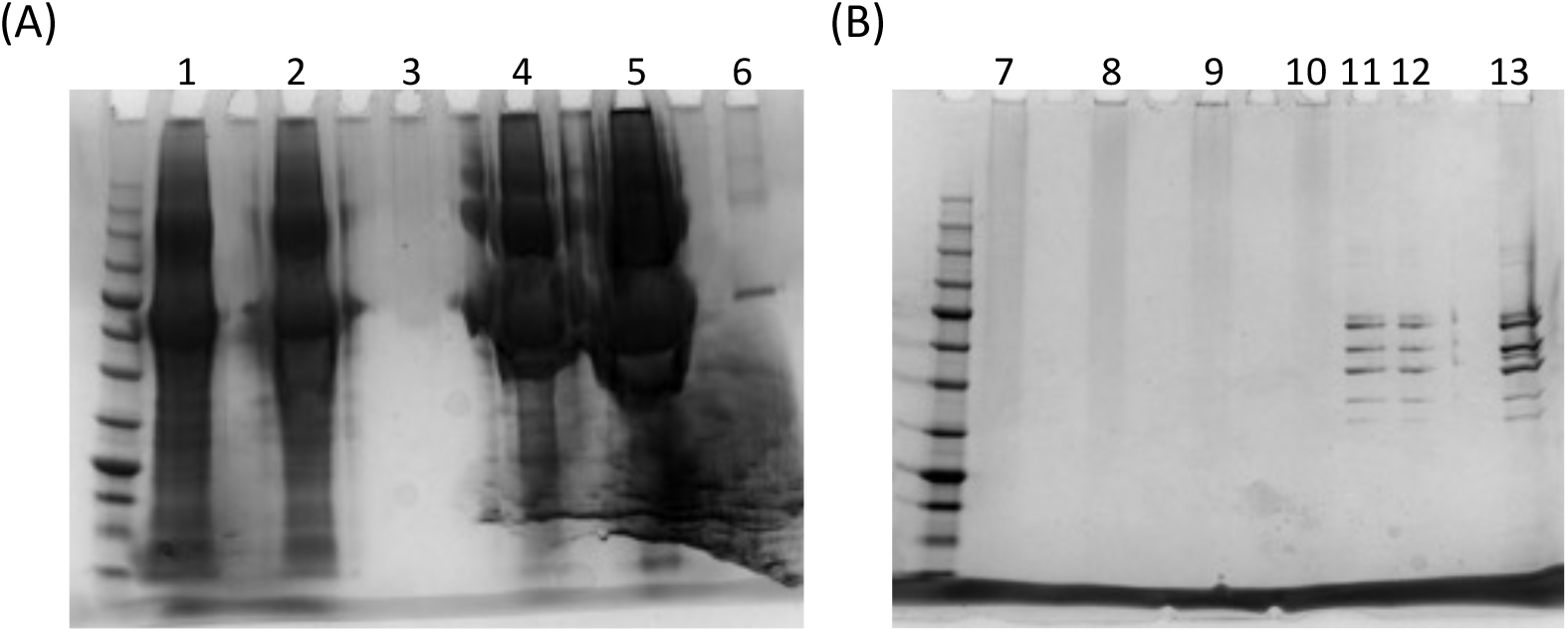
Method development for extracellular protein extraction. Protein extraction of either **(A)** bovine serum albumin (BSA) or **(B)** extracellular proteins collected from *Pseudomonas putida* BIRD-1 spiked into either soil (1,2, 11,12) or water (5, 6, 13) visualised using 1D-SDS PAGE analysis. No spikes controls for either substrate were used as a comparison (3, 7-10).

**Figure S2.**
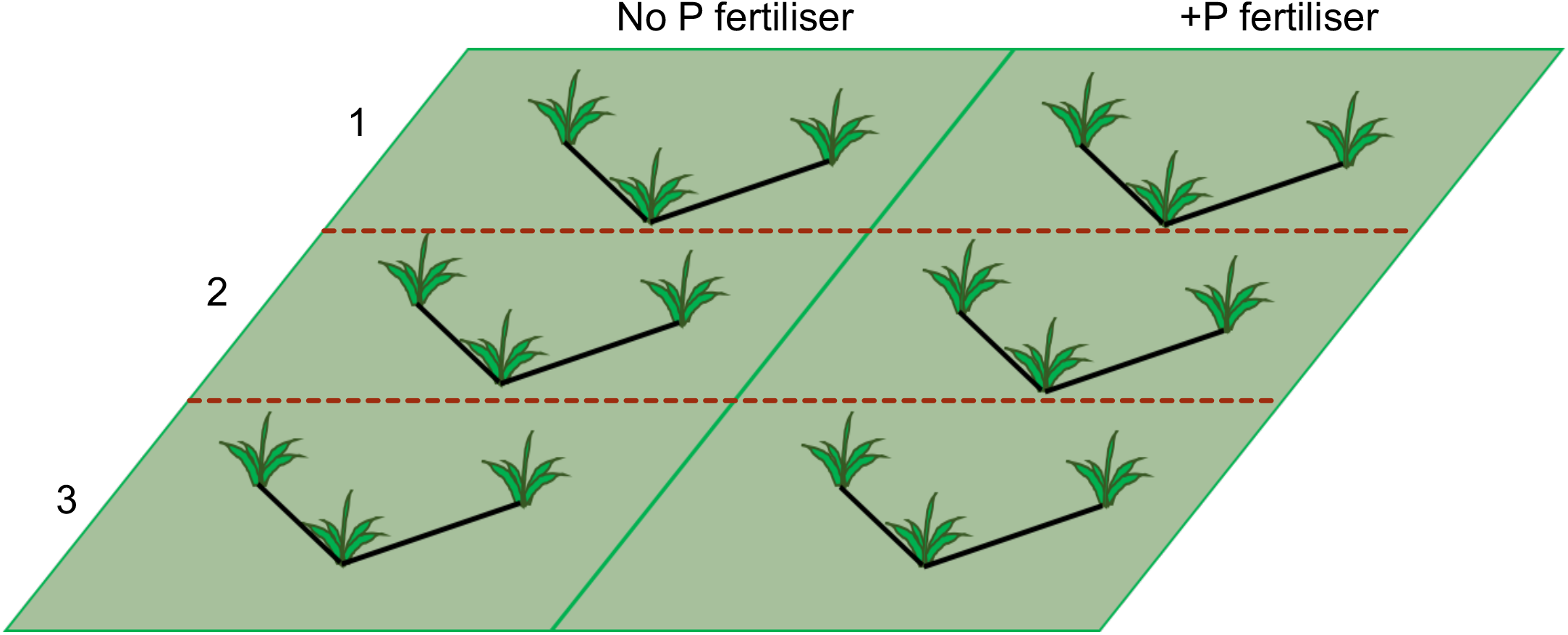
Experimental design of the *Brassica napus* L. field trial. Plants were grown under contrasting fertiliser regimes with different quantities of calcium phosphate applied. Each treatment was divided into three sections and plants were harvested at three distinct sites within the plot. Each ‘V’ was combined to collected 10 g of rhizosphere, in total.

**Figure S3.**
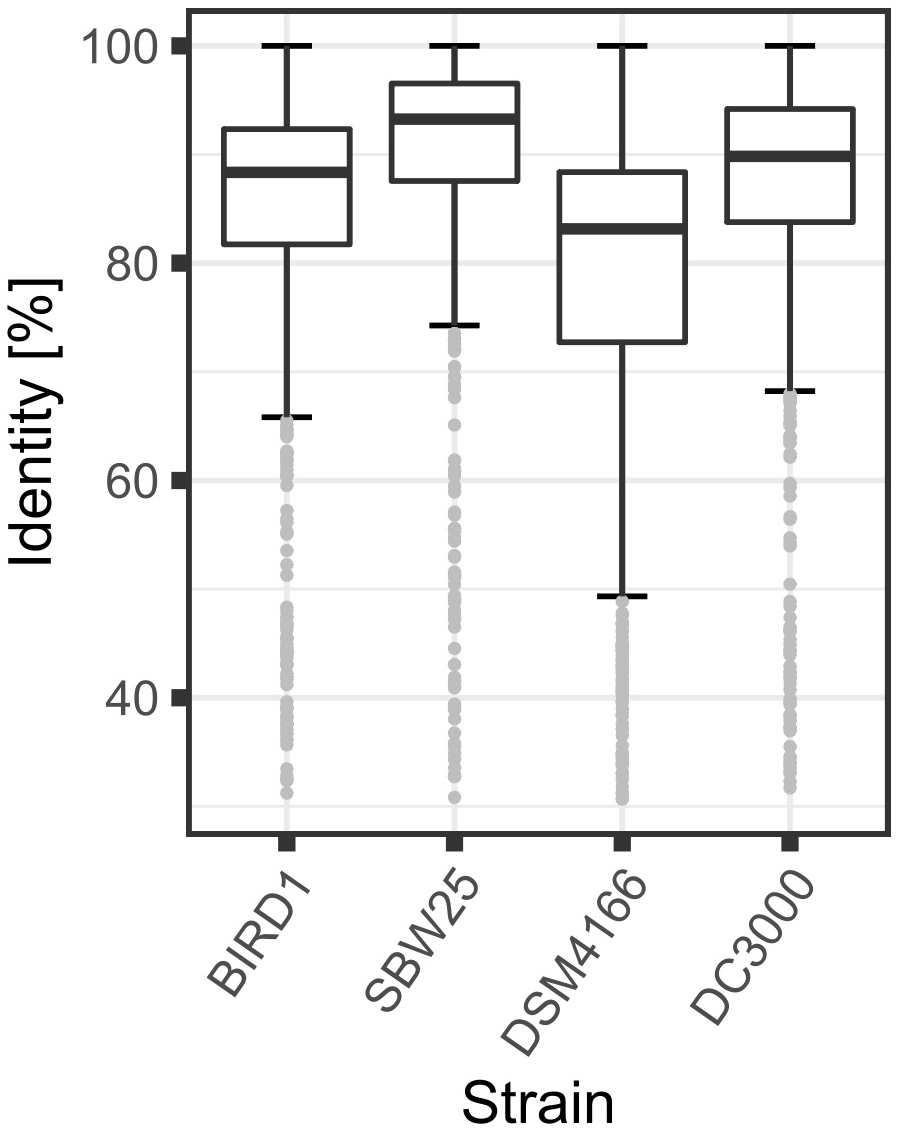
Percentage identity of the *Pseudomonas* related proteins detected in the metaproteome against four *Pseudomonas* ssp. BIRD1, *Pseudomonas putida* BIRD-1; SBW25, *Pseudomonas fluorescens* SBW25; DSM4166, *Pseudomonas stutzeri* DSM4166; *Pseudomonas syringae* DC3000

**Figure S4.**
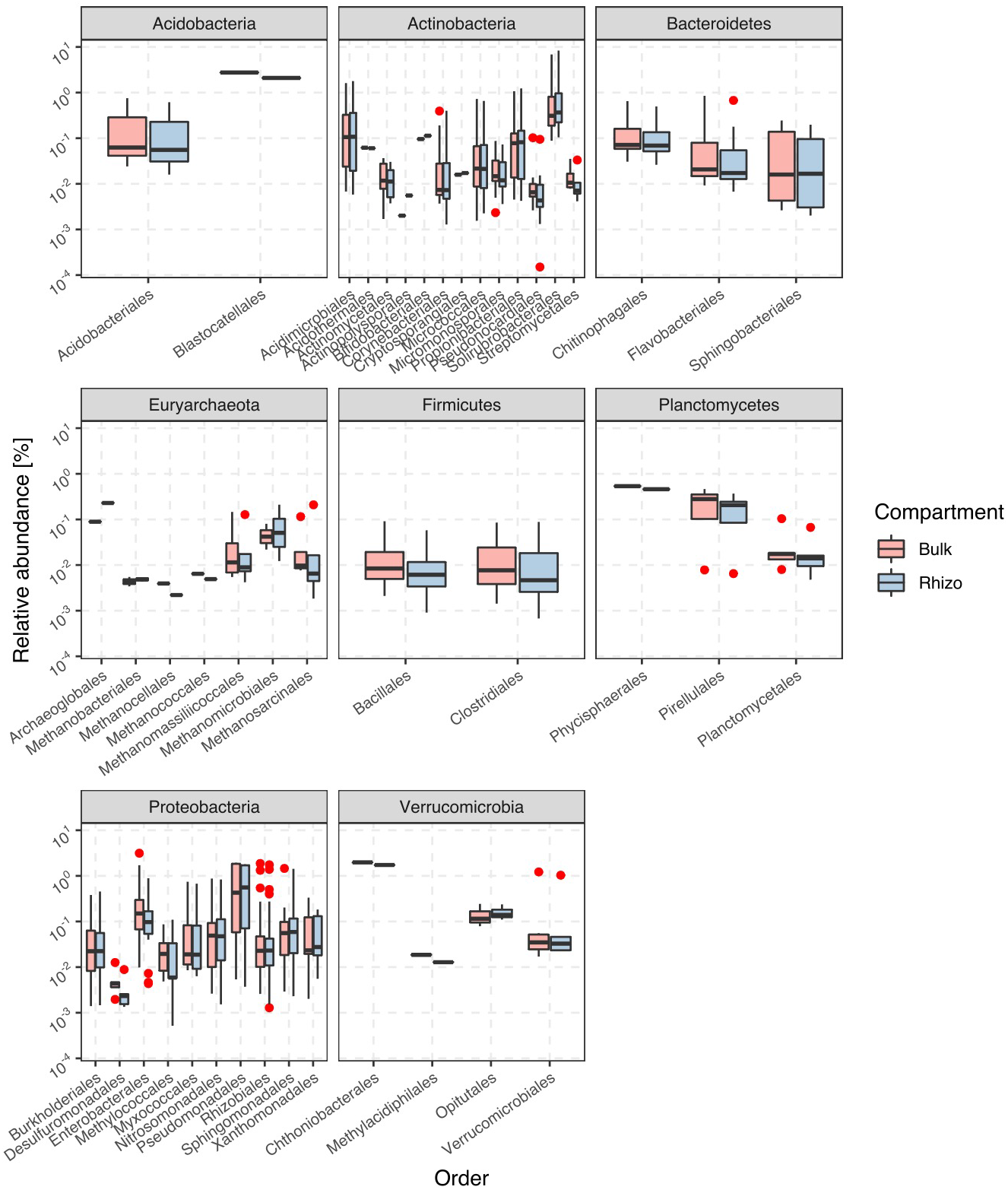
Microbial community associated with field-grown *Brassica napus*. Comparison of bacterial groups at the order-level (separated by phyla) found in the *B. napus* rhizosphere (n=6 MG) compared to the surrounding bulk soil (n= 6 MG). Red data points denote outliers.

**Figure S5.**
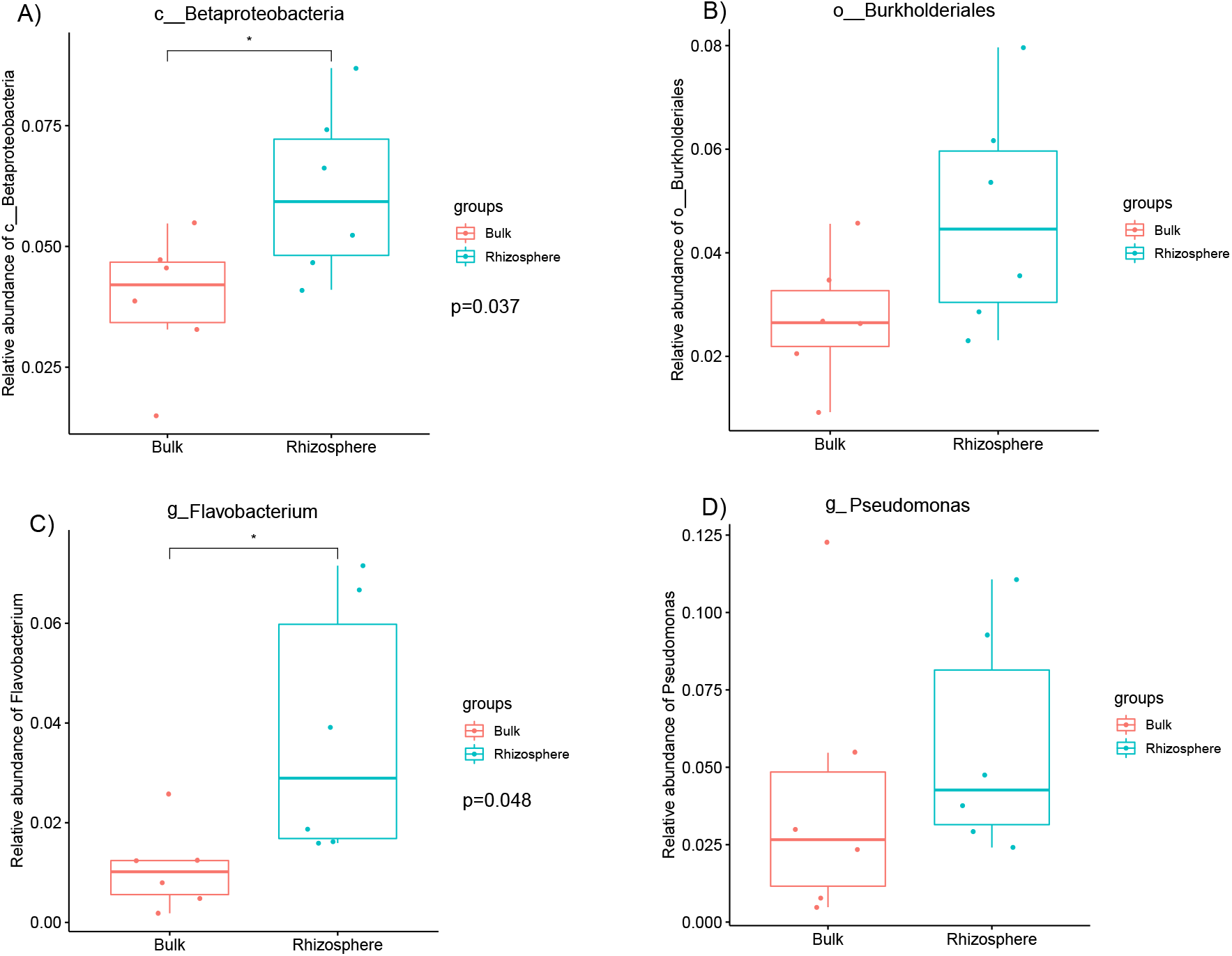
Microbial community associated with field-grown *Brassica napus*. Comparison of bacterial groups at the order-level (separated by phyla) found in the *B. napus* rhizosphere (n=6 MG) compared to the surrounding bulk soil (n= 6 MG). Red data points denote outliers.

**Figure S6.**
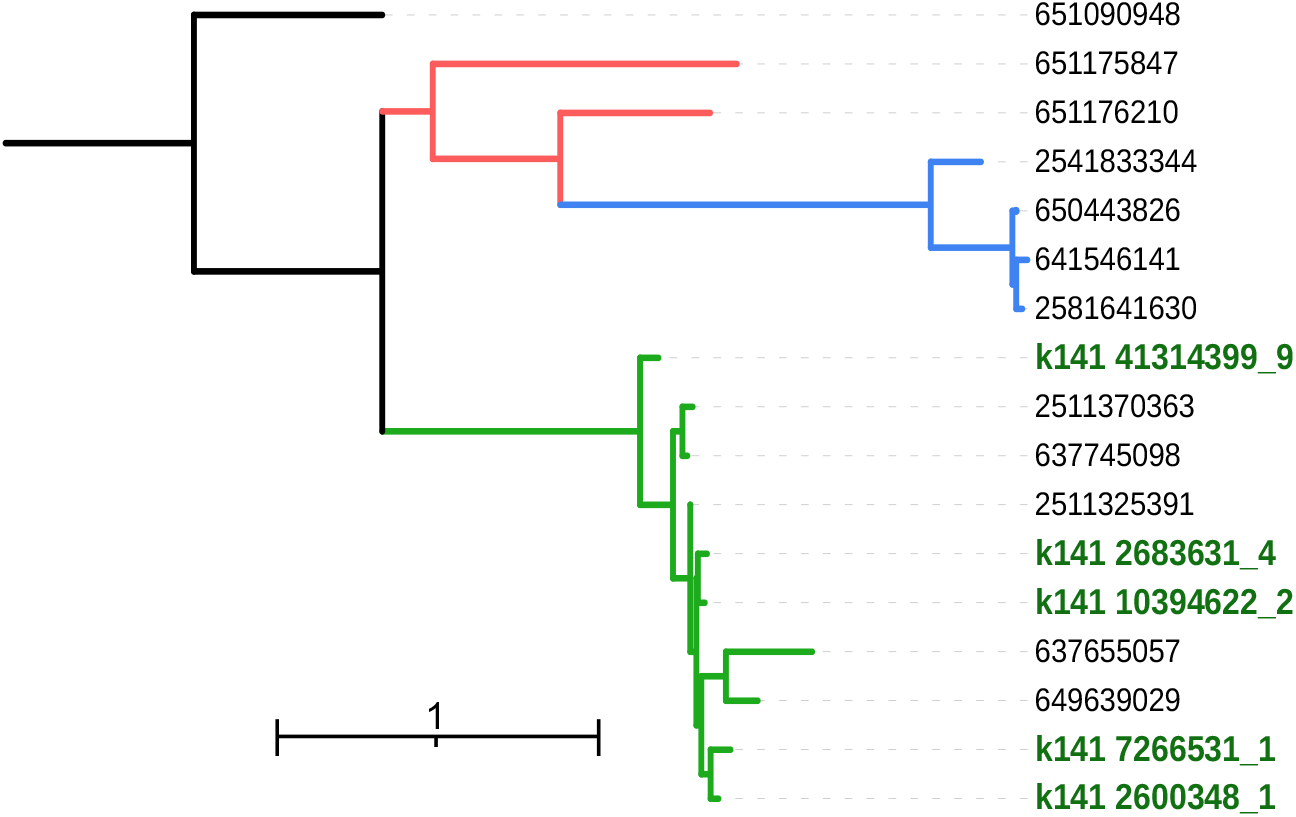
Diversity of PhoX phosphatases detected in the rhizosphere MP. Red denotes *P. stutzeri* PhoX; Blue denotes *P. putida* PhoX; Green denotes *P. fluorescens* PhoX. IMG accession number are denoted in black. ORF numbers for soil PhoX are denoted in green.

**Figure S7.**
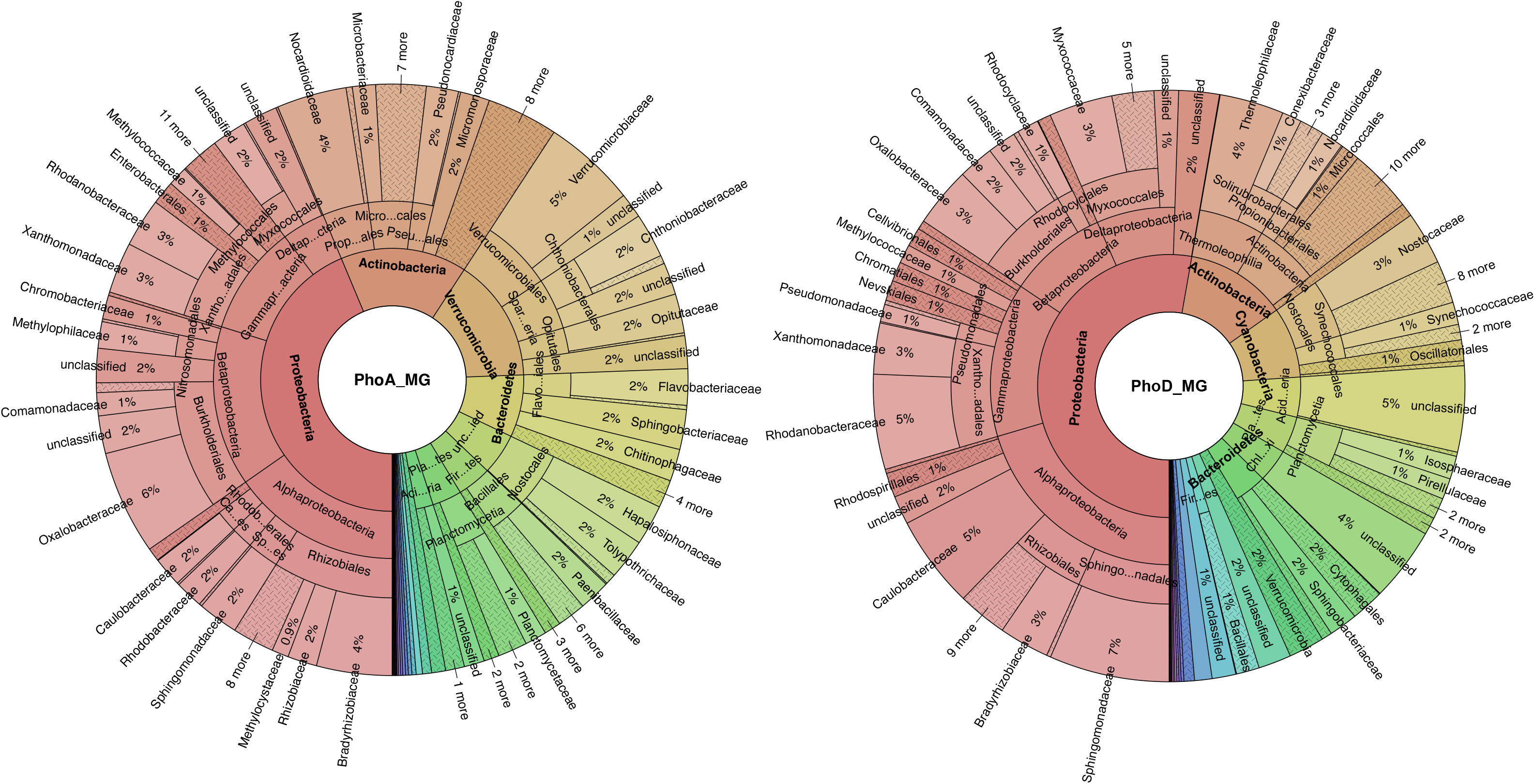
Diversity of alkaline phosphatases detected in the total microbial community (metagenone). Relative abundance, based on normalised read coverage for PhoA **(A)** PhoD **(B)** across both compartments are shown.

